# Prediction of Biological Age and Blood Biomarkers from DNA Methylation Profiles Measured by the Methylation Screening Array: Development and Validation of Models on Japanese Data

**DOI:** 10.64898/2026.02.06.703638

**Authors:** Tatsuma Shoji, Yui Tomo, Ryo Nakaki

## Abstract

**Background:** Epigenetic clocks based on DNA methylation (DNAm) are widely used indicators of biological aging; however, most established models have been developed using EPIC arrays and non-Japanese populations. The Methylation Screening Array (MSA), a cost-efficient platform with reduced CpG content, has not been evaluated for its capacity to support biological age estimation and biomarker prediction in Japanese cohorts.

**Methods:** DNAm profiles and clinical laboratory measurements were obtained from 166 Japanese participants for model development; an independent cohort of 48 individuals processed at a separate institute was used for validation. A linear regression model was trained using the Elastic Net method to predict phenotypic age from MSA-derived methylation data, and a two-stage modeling (residual learning) framework integrating EPIC-based clock predictions with MSA-specific residual predictions was evaluated. Additional models were constructed to examine the predictability of 59 clinical biomarkers and their log-transformed variants, including sex-stratified analyses.

**Results:** The MSA-based model accurately predicted phenotypic age in the validation dataset; prediction performance improved when the EPIC-based estimates were incorporated through the residual learning framework. Several clinical biomarkers, particularly those related to leukocyte composition and sex hormone regulation, were also predicted from the MSA data, although some markers were strongly affected by sex. Some of the nine constituent phenotypic age biomarkers were not individually predicted.

**Conclusions:** MSA methylation profiles contain sufficient biological information for reliable prediction of epigenetic aging markers in Japanese individuals. These findings demonstrate the feasibility of applying cost-efficient MSA-based DNAm profiling for biological age prediction and provide a methodological foundation for expanding epigenetic biomarker applications in Japan.

## Introduction

DNA methylation (DNAm)-based biomarkers of aging, commonly referred to as epigenetic clocks, are highly reproducible predictors of chronological and biological ages in diverse tissues and populations (Horvath, 2013; Hannum et al., 2013; Field et al., 2018; Bell et al., 2019). Early epigenetic clock models established that DNAm patterns change in a highly structured manner with age (Johansson et al., 2013). Horvath’s 2013 cross-tissue clock demonstrated that age-associated methylation signatures are conserved across > 50 tissue and cell types, allowing accurate chronological age estimation using a single predictive model trained on over 8,000 samples (Horvath, 2013). In contrast, the blood-specific clock developed by Hannum et al. in the same year captured hematopoietic and immune system alterations characteristic of aging, correlating strongly with chronological age and mortality risk (Hannum et al., 2013). These foundational models demonstrate that DNAm provides a quantitative molecular framework for studying human aging and motivate the development of clocks that directly reflect biological decline (Horvath & Raj, 2018).

Levine et al. introduced PhenoAge, a model designed to quantify biological age by predicting a composite clinical metric termed phenotypic age (Levine et al., 2018). Phenotypic age was derived from nine blood-based biomarkers: albumin, creatinine, blood glucose, C-reactive protein, lymphocyte percentage, mean cell volume (MCV), red cell distribution width-coefficient of variation (RDW-CV), alkaline phosphatase (ALP), and white blood cell (WBC) counts; these markers collectively capture inflammation, metabolic status, renal function, and hematologic variability, each identified as a predictor of 10-year mortality. After generating this clinical phenotype, Levine et al. constructed a methylation-based model that maps CpG methylation signatures to phenotypic age, resulting in a clock that robustly predicts mortality, morbidity, functional decline, and multiple age-related pathologies (Levine et al., 2018).

A subsequent advancement was the development of GrimAge, which estimates time-to-death by integrating DNAm-derived surrogates of aging-related plasma proteins with a methylation-based estimator of smoking pack-years (Lu et al., 2019). These surrogates reflect pathways related to inflammation, cardiovascular aging, metabolic dysregulation, and systemic physiological decline (Lu et al., 2019; Hillary et al., 2020). GrimAge incorporates these molecular predictors with chronological age and sex to identify biomarkers that are highly predictive of mortality risk. A later refinement, GrimAge v2, expanded the set of protein surrogates and improved adjustment for smoking-associated methylation signals, enhancing performance in younger adults, non-smokers, and diverse ethnic backgrounds (Lu et al., 2021). Together, these clocks illustrate the progressive evolution of epigenetic aging metrics toward greater biological relevance and predictive accuracy (Lu et al., 2019).

Despite rapid advancement, most established epigenetic clocks, including the Horvath, Hannum, PhenoAge, and GrimAge clocks, have been constructed primarily using data from North American and European populations, raising concerns about their applicability to individuals with distinct genetic backgrounds, environmental exposures, and lifestyle patterns (Horvath, 2013; Hannum et al., 2013; Levine et al., 2018; Lu et al., 2019). Recent studies have highlighted the need for population-specific epigenetic aging models. An example is the Japanese PCPhenoAge (J-PCPhenoAge), which uses a principal component-based framework and a transfer learning approach to improve biological age estimation in Japanese cohorts (Tomo et al., 2024). However, J-PCPhenoAge and most other epigenetic clocks have been developed using Infinium EPIC or EPIC v2 methylation arrays, whose CpG coverage differs substantially from that of the newer and more cost-effective Methylation Screening Array (MSA). Since approximately half of the CpG sites in the MSA platform are not present on the EPIC-based platform, it remains unclear whether EPIC-based epigenetic aging metrics can be accurately reproduced using MSA data. This discrepancy constitutes a methodological gap, particularly in Japan, where MSA is increasingly adopted in clinical and research applications. An MSA-compatible Japanese biological age model that can reliably predict phenotypic age is required.

Beyond age estimation, DNAm profiles also encode information related to circulating clinical biomarkers, enabling indirect physiological state inference from epigenomic data. DNAm signatures can predict blood-based traits, such as inflammatory markers, immune cell composition, lipid profiles, and hormone levels, suggesting that methylation captures integrative signals reflecting genetic regulation and environmental exposure (Houseman et al., 2012; Lu et al., 2019; Hillary et al., 2020; McCartney et al., 2021). These DNAm-derived biomarker proxies are associated with disease risk, functional decline, and mortality (Marioni et al., 2015; Levine et al., 2018; Lu et al., 2019; Hillary et al., 2020; McCartney et al., 2021), highlighting their potential utility in large-scale epidemiological and clinical studies where direct biomarker measurements may be unavailable or impractical. However, most existing biomarker prediction models were developed using EPIC or earlier high-density arrays, leaving the applicability of such approaches to lower-density platforms largely unexplored. Given that the MSA was specifically designed to enrich CpG sites relevant to aging, chronic disease, immune function, and metabolic regulation, it represents a promising platform for biomarker inference. However, a systematic evaluation of whether blood-based biomarkers can be reliably predicted from MSA-derived methylation data is lacking. Establishing MSA-based biomarker prediction models is essential to extend the clinical and translational utility of cost-efficient methylation profiling, particularly in population-specific contexts such as Japanese cohorts.

This study aimed to develop a method to evaluate biological age for the Japanese population from DNAm profiles measured by MSA, by constructing an MSA-based phenotypic age prediction model and validating its performance in independent Japanese cohorts. We collected MSA methylation profiles and clinical laboratory data from 166 individuals to train an elastic net-based predictor of phenotypic age, followed by validation using an additional cohort of 48 individuals processed at a separate institute. Next, we examine a two-stage modeling strategy that integrates EPIC-based clock estimates with MSA-specific residual predictions to improve cross-platform compatibility. Finally, we evaluated whether individual clinical biomarkers, including the nine components used in the phenotypic age calculation, could be predicted from DNAm profiles to assess its utility in clinical biomarker evaluation.

## Materials & Methods

### Study Participants and Sample Collection

Whole-blood samples were obtained according to the protocol described by Tomo et al. (2025). Briefly, blood samples were collected from healthy adult volunteers recruited in Japan using two independently approved observational research protocols. A total of 214 unique individuals participated, divided into 166 in the training dataset and 48 in the validation dataset. A total of 166 samples were collected at Y’s Science Clinic Hiroo (Minato-ku, Tokyo, Japan) as part of the project, “Evaluation of Biological Age Based on DNA Methylation and Its Clinical Significance,” which was reviewed and approved by the Shiba Palace Clinic’s Institutional Review Board. Written informed consent was obtained from all participants. An independent validation cohort of 48 individuals was collected under the project “Evaluation of Epigenetic Clock Measurement in Clinical Practice,” approved by the Asia–Oceania Anti-Aging Promotion Association Ethics Review Committee (AOAAPA22-001, approved on May 27, 2022). Genomic DNA was extracted using a Maxwell RSC Blood DNA Kit (Promega, Madison, WI, USA) according to the manufacturer’s instructions. All procedures adhered to the Ethical Guidelines for Medical and Biological Research Involving Human Subjects issued by the Japanese Government.

### Clinical Laboratory Measurements

A total of 59 clinical laboratory markers were measured in whole blood samples collected at the time of DNA extraction, including standard biochemical markers, hematological indices, hormone measurements, inflammatory markers, lipid parameters, and protein fractionation profiles. The assessed items were ALP based on International Federation of Clinical Chemistry and Laboratory Medicine (ALP/IFCC), alanine aminotransferase (ALT), glutamic pyruvate transaminase (GPT), aspartate aminotransferase (AST), glutamic oxaloacetic transaminase, quantitative C-reactive protein, calcium, chloride, dehydroepiandrosterone sulfate (DHEA-S), follicle-stimulating hormone (FSH), serum iron, hematocrit, high-density lipoprotein (HDL) cholesterol, hemoglobin (Hb), HbA1c, potassium, lactate dehydrogenase based on IFCC (LD/IFCC), low-density lipoprotein cholesterol, luteinizing hormone (LH), mean corpuscular Hb (MCH), MCH concentration, mean corpuscular volume (MCV), magnesium, sodium, N-terminal pro-brain natriuretic peptide (NT-proBNP), phosphorus, red blood cell (RBC) count, RDW-CV, thyroid-stimulating hormone, WBC count, γ-glutamyl transpeptidase (γ-GTP), albumin, insulin, estradiol, creatinine, total testosterone, free triiodothyronine (free T3) and thyroxine (free T4), progesterone, triglycerides, blood urea nitrogen, uric acid, differential leukocyte counts (lymphocytes, monocytes, neutrophils, basophils, and eosinophils), total cholesterol, total bilirubin, total protein, α1-globulin, α2-globulin, β-globulin, and γ-globulin, albumin/globulin ratio (A/G ratio), albumin fraction, platelet count, serum amylase, blood glucose, free testosterone, and high-sensitivity prostate-specific antigen (PSA). All measurements were performed in two analytical batches.

### Outlier Detection and Missing-Value Imputation

Outlier detection was performed for each clinical laboratory marker using the conditional residual approach. A Huber regression model with epsilon = 1.35 was fitted using the remaining 58 markers as predictors, and residuals were standardized using the median absolute deviation. Values with a standardized conditional residual ≥ 6 were identified as potential outliers and flagged for exclusion. In addition, samples exhibiting statistically significant Mahalanobis distances, as determined by a chi-square test at a significance level of 0.0001 and computed from the standardized feature space, were considered for removal. No individual measurements or samples met the exclusion criteria. The proportions of missing values for each marker are summarized in Table S1. Missing data were imputed using a multivariate single imputation strategy based on the ExtraTrees algorithm (Geurts et al., 2006). The imputed values for all 214 participants are provided in Table S2. Outlier detection and missing-value imputation were performed using Scikit-learn (version 1.7.1) in Python (version 3.12.3).

### Calculation of Phenotypic Age

Phenotypic age was calculated following the procedure described in the original PhenoAge publication, which derived a mortality-optimized biological age metric based on nine clinical biomarkers (Levine et al., 2018). Albumin values were computed from the total protein and albumin fractions obtained through serum protein fractionation, whereas the remaining biomarkers were converted to the units required by the PhenoAge algorithm. After harmonizing measurement units, the nine biomarkers—albumin (g/L), creatinine (µmol/L), blood glucose (mmol/L), C-reactive protein (mg/L), lymphocyte percentage (%), MCV (fL), RDW-CV (%), ALP (U/L), and WBC count (1000 cells/µL)—were incorporated into the published mortality-score equation. Specifically, mortality score (MS) was calculated as follows:

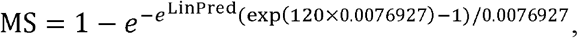

where

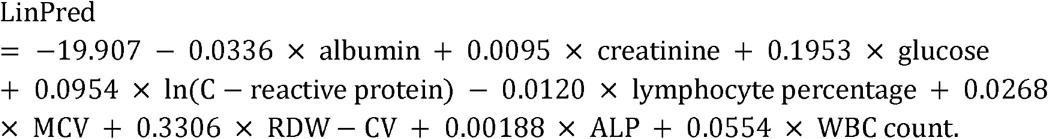

Phenotypic age was then calculated by transforming this mortality score according to the following equation:

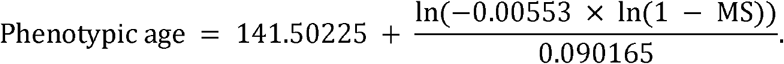

All calculations were performed using Python (version 3.12.3) according to the specifications described in the original method.

### DNA Methylation Profiling and Preprocessing

Genome-wide DNAm profiling was performed partially according to the same protocol as Tomo et al. (2025). Briefly, DNAm profiling was performed using the Illumina Infinium MSA. Array processing and scanning for a training dataset consisting of 166 samples were conducted by Rhelixa Inc. (Chuo-ku, Tokyo, Japan); the validation dataset of 48 samples was processed independently by CD Genomics (Shirley, NY, USA). No individuals overlapped between the two datasets. After bisulfite conversion according to Illumina’s recommended protocol, the converted DNA was amplified, fragmented, hybridized to the array, and scanned using an Illumina iScan System. Raw IDAT files were processed in R (version 4.4.2) using the SeSAMe pipeline (version 1.24.0) (Triche et al., 2013; Zhou et al., 2018), including background correction with the normal-exponential out-of-band (noob) method and application of the pOOBAH detection mask for probe-level quality filtering. Beta values were calculated as the ratio of methylated signal intensity to the sum of methylated and unmethylated signal intensities (Aryee et al., 2014). Probes failing the detection thresholds, overlapping single-nucleotide polymorphisms likely to affect hybridization, exhibiting cross-reactive mapping to multiple genomic loci, and located on sex chromosomes were removed before analysis to ensure high-quality data. Beta value matrices are available upon request.

### Computation of Existing Epigenetic Clocks

Four established epigenetic clocks (Horvath, Hannum, PhenoAge, and GrimAge) were calculated using DNAm data from MSA. Each indicator was computed according to published algorithms and corresponding regression coefficients. The Horvath and Hannum clocks were derived using CpG sets and coefficients from their original publications (Horvath, 2013; Hannum et al., 2013). PhenoAge was calculated according to the procedure described by Levine et al. (2018), which maps methylation values to phenotypic age. GrimAge estimation was performed according to the model introduced by Lu et al. (2019), which integrates DNAm-derived surrogates of age-related plasma proteins with a DNAm-based smoking exposure estimator. Additionally, we derived J-PhenoAge, a Japanese PhenoAge, based on the EPICv2 array. The J-PhenoAge model was specified as a linear regression model including 3,586 candidate CpGs as features, and trained using data from 143 Japanese individuals collected by Tomo et al. (2024) and the Transfer Elastic Net method following the same procedures, except for principal component transformation. The candidate CpGs were selected to comprehensively include sites incorporated into previously published epigenetic clocks. Subjects were collected under the project “Evaluation of Epigenetic Clock Measurement in Clinical Practice,” approved by the Asia-Oceania Anti-Aging Promotion Association Ethics Review Committee (approval number AOAAPA22-001, approved on May 27, 2022); none of these subjects were included in the 48-participant validation dataset described above. The estimated coefficients for J-PhenoAge are listed in Table S3. We then computed J-PhenoAge using the DNAm data from the MSA.

### Construction of the MSA-based Phenotypic Age Model

To develop an MSA-based phenotypic age model, a linear regression model was constructed using CpG beta values as predictors and phenotypic age as the target variable, trained using the Elastic Net method (Zou and Hastie, 2005). All features were standardized before training. The dataset of 166 participants was randomly divided such that 80% of the samples were used for hyperparameter tuning with fivefold cross-validation, whereas the remaining 20% served as the internal test set. After the optimal hyperparameters were identified, the final model was evaluated on a held-out 20% subset by computing the correlation coefficient, mean absolute error (MAE), and root mean squared error (RMSE) between predicted and observed phenotypic age. Validation was then performed using an independent dataset of 48 participants processed at a separate institute, and the same performance metrics were computed. CpG sites and regression coefficients used in the final model are summarized in Table S4. All calculations and visualizations were performed using Matplotlib (version 3.10.6) and Scikit-learn (version 1.7.1) in Python (version 3.12.3).

### Two-Stage Prediction Model

A two-stage modeling framework was implemented to evaluate whether predictive performance could be further improved by leveraging information from EPIC-based clocks. We applied a residual learning framework, in which the residuals from regressing phenotypic age on existing epigenetic clock predictions were used as targets for training on the MSA-measured CpG sites. This can be regarded as a transfer learning approach, in which knowledge from established epigenetic clocks is refined using a new set of epigenetic features (Kuzborskij and Orabona, 2013; Zheng et al., 2021).

In the first stage, five epigenetic clocks (Horvath, Hannum, PhenoAge, GrimAge, and J-PhenoAge) were computed for all 166 participants; a linear regression model was trained using ℓ_2_ regularization (Ridge regression) to predict phenotypic age using these five standardized clock values as predictors. In the second stage, residuals obtained by subtracting the first-stage predictions from the observed phenotypic ages were used as target variables; MSA CpG beta values were used as predictors in a new linear regression model and trained using the Elastic Net method. For validation, 48 independent samples were used to generate the first-stage predicted phenotypic age values and second-stage predicted residuals, which were then summed to obtain final predictions. Model performance was evaluated by computing the correlation coefficient, MAE, and RMSE between the predicted and observed phenotypic ages. The coefficients used in the first- and second-stage models are summarized in Tables S5 and S6. All calculations and visualizations were performed using Matplotlib (version 3.10.6) and Scikit-learn (version 1.7.1) in Python (version 3.12.3).

### Prediction of Clinical Biomarkers from MSA Data

Predictive models were constructed for all 59 laboratory measurements, as well as the log-transformed versions of six variables exhibiting right-skewed distributions (γ-GTP, NT-proBNP, triglycerides, AST, ALT, and quantitative C-reactive protein), to assess whether individual clinical biomarkers could be inferred directly from MSA-derived methylation data. In Model 1, a linear regression model using CpG beta values as predictors and each biomarker as the target was specified and trained using the Elastic Net method, following the same training, testing, and validation framework used for the phenotypic age model. Performance was evaluated in the validation cohort of 48 individuals by computing the correlation coefficient, MAE, and RMSE between observed and predicted values. In addition, partial correlation coefficients controlling for sex were calculated for all biomarkers to account for potential confounding factors. In Model 2, participants were stratified by sex, and separate models were trained for males and females using CpG beta values and chronological age as predictors; performance was evaluated for each sex using the validation cohort based on correlation, MAE, and RMSE. CpG sites and regression coefficients used in the final models are available upon request. MAE and RMSE values are summarized in Tables S3 and S4, respectively. All calculations and visualizations were performed using Matplotlib (version 3.10.6) and Scikit-learn (version 1.7.1) in Python (version 3.12.3).

## Results

### Prediction of Phenotypic Age Using MSA Data

In the validation cohort of 48 individuals, the predicted phenotypic ages showed strong correlations with observed values (R=0.930, MAE=5.565, RMSE=7.462) (Fig. 1). These results suggest that phenotypic age could be accurately predicted from MSA methylation profiles.

**Figure 1.**
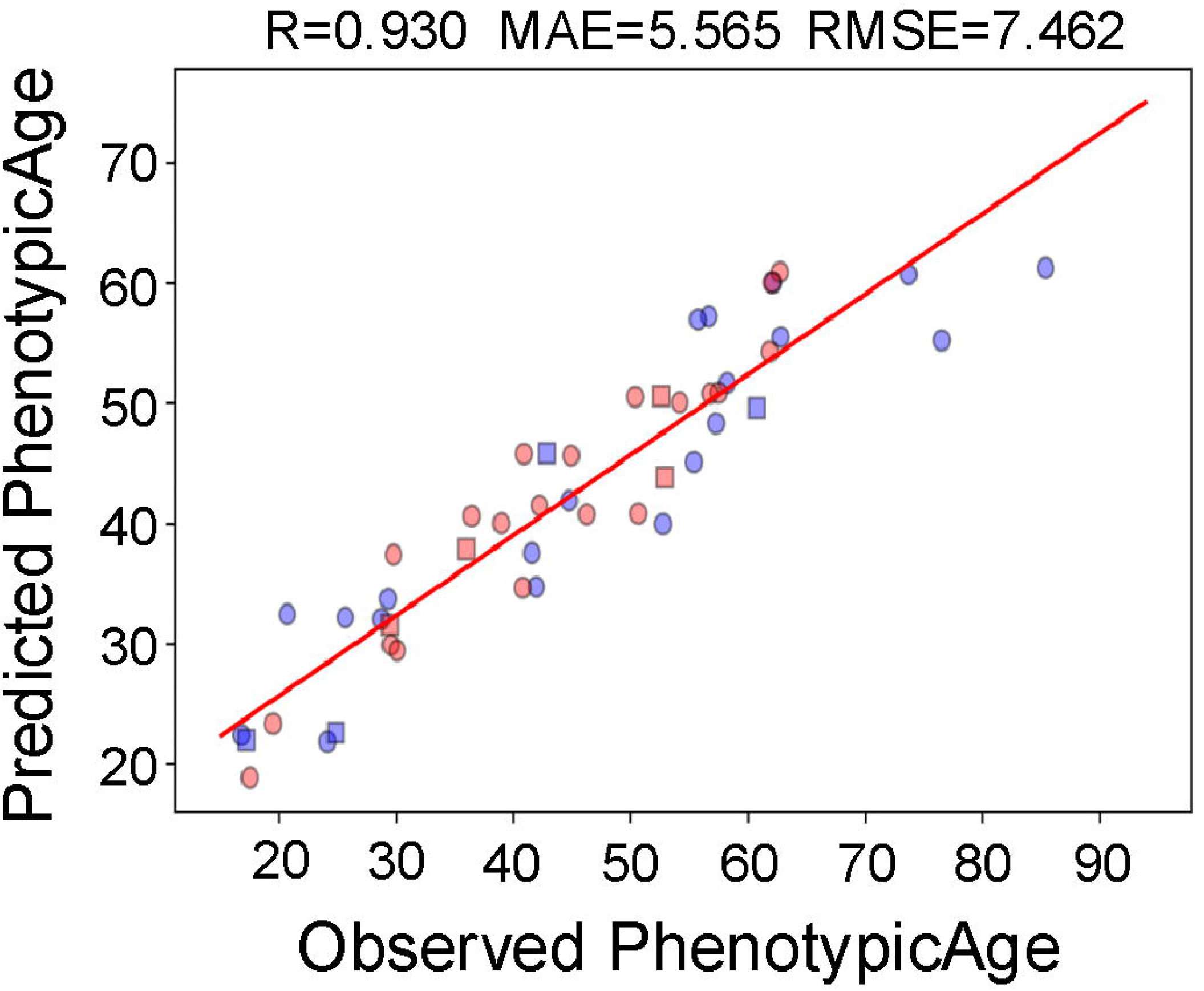
Scatter plot comparing observed and predicted phenotypic age in the validation cohort. Scatter plot of observed phenotypic age values versus predictions generated by the MSA-based phenotypic age model for 48 validation samples. Blue points represent male participants, and red points represent female participants. Squares indicate samples processed in batch 1, whereas circular markers represent samples processed in batch 2. The correlation coefficients MAE and RMSE are presented at the top of the panel.

### Two-Stage Prediction Model

To assess whether the predictive accuracy could be enhanced by integrating EPIC-based epigenetic aging information with MSA-specific methylation signals, a two-stage prediction framework was evaluated. In the validation cohort of 48 individuals, predicted residuals correlated positively with observed residuals (R=0.277) (Fig. 2A). The correlation coefficient increased when the first- and second-stage predicted residuals were combined to generate the final phenotypic age predictions, and both MAE and RMSE decreased relative to the single-stage model (R=0.945, MAE=4.192, RMSE=5.509) (Fig. 2B).

**Figure 2.**
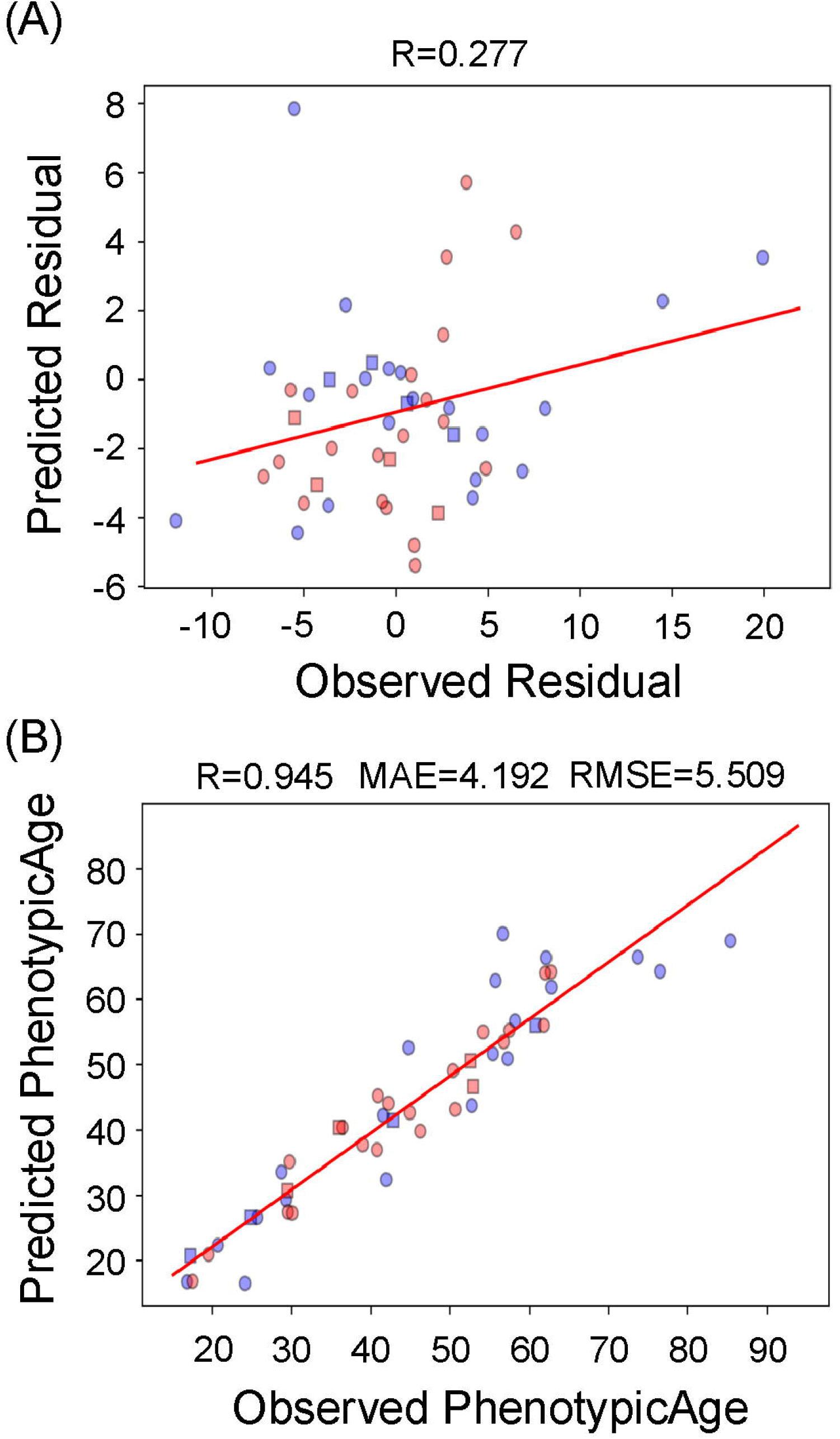
Scatter plots illustrating the two-stage prediction framework in the validation cohort. **(A)** Scatter plot comparing observed and predicted residuals obtained from the first-stage model for the 48 validation samples. The correlation coefficient is shown at the top of the panel. **(B)** Scatter plot comparing the observed and predicted sums of the first-stage estimates and second-stage predicted residuals. The correlation coefficient, MAE, and RMSE are shown at the top of the panel. Blue points represent male participants and red points represent female participants. Square markers indicate samples processed in batch 1, whereas circular markers represent samples processed in batch 2.

### Prediction of Clinical Biomarkers from MSA Data

Expanding upon successful phenotypic age prediction, we examined whether individual clinical biomarkers could be inferred directly from MSA-derived methylation profiles. Model 1 demonstrated favorable predictive performance for several hematological indices and sex hormone-related markers in the validation cohort (Fig. 3). However, for certain biomarkers such as creatinine, scatterplots revealed distinct clusters by sex (Fig. 4).

**Figure 3.**
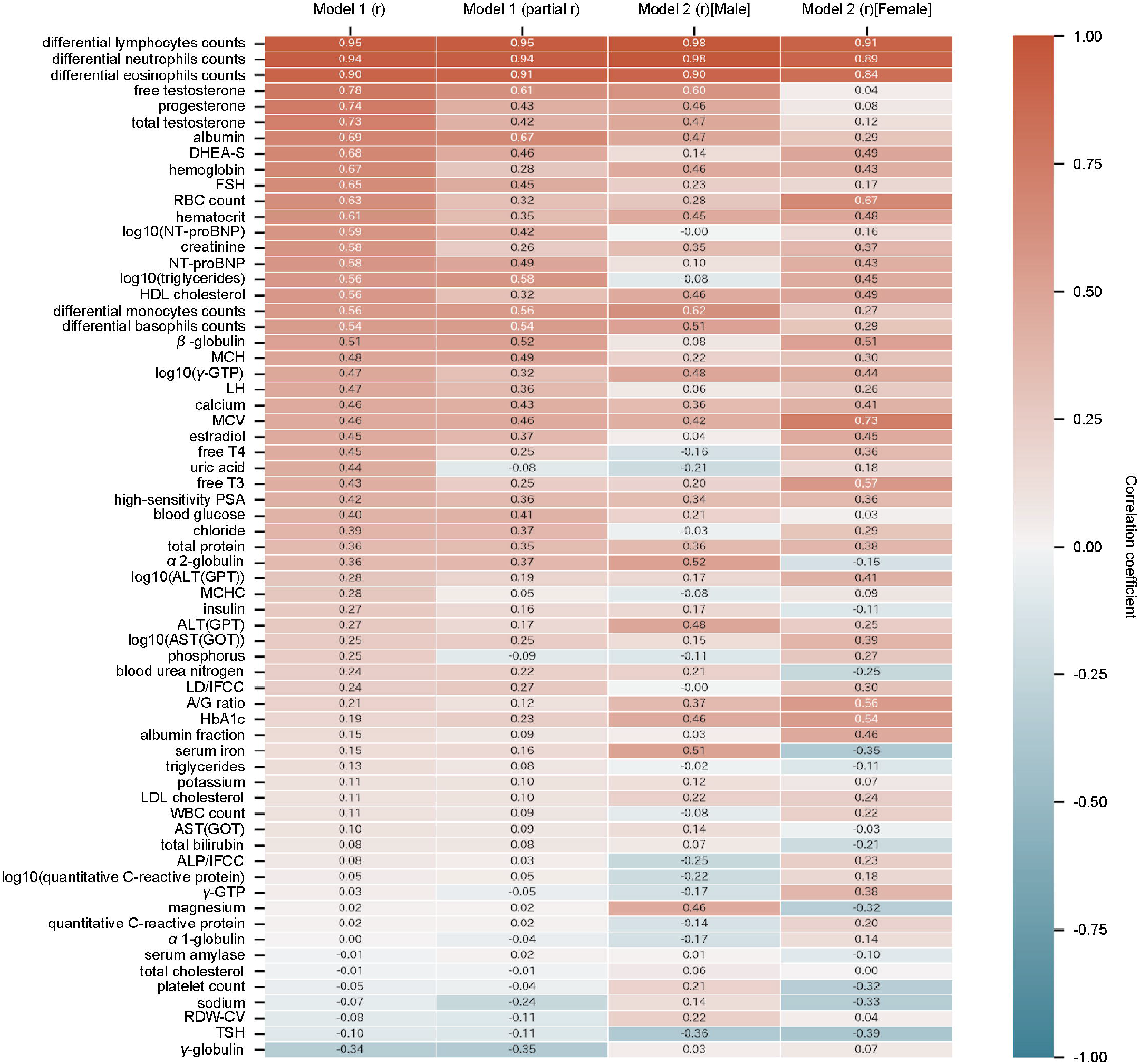
Correlations between observed and predicted clinical biomarkers in the validation cohort. Correlation coefficients between the observed values and predictions for all 59 clinical biomarkers in the validation cohort of 48 individuals. Model 1 shows the correlation across all participants (Model 1 (r)) and the sex-adjusted partial correlation (Model 1 (partial r)). Model 2 presents the sex-stratified prediction performance, with separate correlation coefficients for males (Model 2 (r) [Male]) and females (Model (r) [Female]).

**Figure 4.**
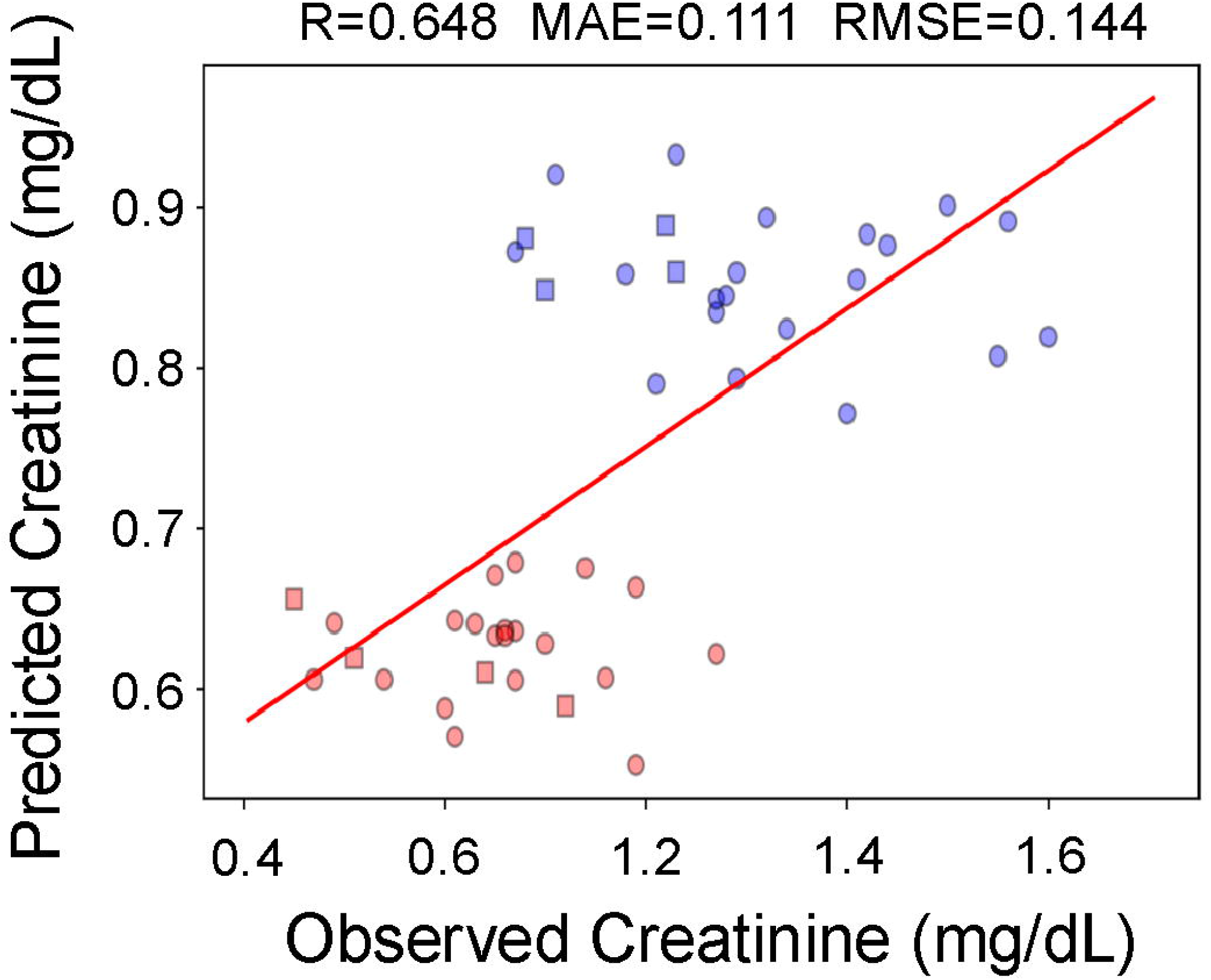
Scatter plot comparing observed and predicted creatinine levels in the validation cohort. Scatter plot of observed creatinine values versus Model 1 predictions for 48 validation samples. Blue points represent male participants, and red points represent female participants. Squares indicate samples processed in batch 1, whereas circular markers represent samples processed in batch 2. The correlation coefficients MAE and RMSE are presented at the top of the panel.

Partial correlation coefficients were then calculated while controlling for sex. Multiple markers, including creatinine and uric acid levels, exhibited reduced associations with observed and predicted values (Fig. 3). Differential lymphocyte, neutrophil, and eosinophil counts, free testosterone, progesterone, total testosterone, albumin, DHEA-S, Hb, FSH, RBC count, hematocrit, NT-proBNP, triglycerides, HDL cholesterol, differential monocyte counts, differential basophil counts, β-globulin, MCH, γ-GTP, LH, calcium, MCV, estradiol, free T4, free T3, high-sensitivity PSA, and blood glucose were predictable biomarkers (correlation coefficient > 0.4, partial correlation coefficient > 0.3) in Model 1.

Model 2 incorporated sex-stratified training to further evaluate the influence of sex on the prediction accuracy (Fig. 3). This approach yielded heterogeneous outcomes: performance improved for certain analytes (such as HbA1c), remained unchanged for some (such as differential lymphocyte counts), and decreased for others (such as total testosterone). Additionally, some biomarkers showed sex-specific predictability patterns; DHEA-S displayed high predictive accuracy only in females, whereas free testosterone exhibited stronger performance in males. The MAE and RMSE of all biomarkers except total testosterone, DHEA-S, estradiol, LD/IFCC, triglycerides, and WBC count decreased compared to those from Model 1 (Tables S3 and S4). α2-globulin, ALT, GPT, A/G ratio, HbA1c, albumin fraction, serum iron, and magnesium were predictable biomarkers in Model 2.

## Discussion

This study presents the first epigenetic clock constructed using MSA in a Japanese population. MSA-based methylation profiles accurately predicted phenotypic age in validation cohorts, indicating that the MSA platform provides sufficient methylation coverage to develop a Japanese epigenetic clock. Predictive accuracy improved further with two-stage modeling using EPIC-based clocks to generate first-stage predictions, and MSA-specific methylation signals captured residual variance not explained by the clocks. Several clinical biomarkers, particularly hematological parameters and sex hormone-related analytes, could be predicted from MSA methylation profiles, although predictability varied by biomarker and was strongly confounded by sex in some cases. These findings fill a methodological gap by enabling reliable age and biomarker prediction using MSA data, and highlight the broader potential of MSA-based epigenetic analyses in Japanese populations.

The strong performance of the MSA-based Japanese phenotypic age model indicates that the reduced CpG content of the MSA platform does not compromise its ability to capture the core methylation signatures underlying phenotypic age in this population. However, the biological information used to construct the MSA-based model may differ from that of J-PhenoAge because only eight common CpG sites were selected by these models; the numbers of selected CpG sites used in the MSA-based model and J-PhenoAge were 174 and 744, respectively. Although there were few common CpG sites between the MSA-based and EPIC-based models, MSA provided sufficient methylome resolution for constructing reliable biological age predictors, supporting the feasibility of applying the Japanese phenotypic age prediction model in future studies that rely on cost-efficient methylation profiling.

The improvement observed in the two-stage prediction framework highlights the complementary nature of the methylation information captured by the EPIC-based clocks and the MSA platform. Although the first-stage model, which integrates existing epigenetic clocks, accounts for a substantial portion of phenotypic age variance, the correlation between the observed and predicted residuals in the second stage demonstrated that MSA contains additional methylation signals that are not represented in EPIC-based features. This suggests that certain age-associated loci, present in MSA but absent from EPIC arrays, contribute meaningful biological information relevant to aging in Japanese individuals. The enhanced predictive performance obtained by combining first-stage predictions with second-stage residual predictions may reflect the ability of MSA to detect platform-specific methylation patterns that refine biological age prediction. These findings indicate that residual learning serves as an effective strategy for harmonizing information across distinct methylation platforms and illustrate the potential utility of residual transfer learning frameworks in future epigenetic aging research.

The heterogeneous performance observed in clinical biomarker prediction provided important insights into the biological specificity of methylation signatures captured by the MSA platform. Although phenotypic age was predicted with high accuracy, its nine constituent biomarkers were not necessarily well predicted, indicating that the composite weighting scheme used in the formula carries critical information that cannot be recovered from individual biomarker predictions alone (Levine et al., 2018; Liu et al., 2018). In contrast, several other biomarker categories, particularly those related to leukocyte composition and sex hormones, showed favorable predictive performance in Model 1; this was consistent with the MSA design, which prioritizes CpG sites associated with lifestyle-related diseases, inflammation, immune function, and aging phenotypes (Hillary et al., 2020). However, sex strongly influenced the prediction accuracy of several analytes, as evidenced by inflated pooled correlations, which diminished substantially after adjusting for sex or stratifying the analysis. These sex dependencies align with those of previous reports, indicating that epigenetic aging signatures and related physiological markers exhibit sexually dimorphic patterns (McCartney et al., 2019). These results indicate that while a subset of biomarkers can be reliably predicted from MSA methylation profiles, predictability is highly trait-dependent and must be interpreted considering sex-specific biological variation.

This study had some limitations. Despite accurate prediction of phenotypic age as a composite measure, the nine constituent biomarkers were not necessarily individually predictable, suggesting that the formula calculating phenotypic age cannot be applied to CpG data alone. This indicates the need to re-estimate biomarker coefficients in the formula specifically for Japanese populations to more accurately reflect population-specific physiological aging. Future studies should focus on expanding sample sizes, refining sex-specific models, and estimating clinical parameter coefficients underlying phenotypic age to enhance the generalizability and biological validity of epigenetic aging models.

Overall, the present findings demonstrate that MSA-based methylation profiling enables reliable phenotypic age prediction, supporting the broader application of cost-efficient epigenetic biomarkers in Japanese clinical and research settings.

## Conclusions

This study aimed to investigate whether MSA could support reliable prediction of biological age and clinical biomarkers in Japanese individuals by developing an MSA-based version of the phenotypic age prediction model and evaluating its performance using independent datasets. Phenotypic age could be accurately predicted from MSA methylation profiles, with accuracy improving when EPIC-based information was incorporated through a residual learning framework. Several clinical biomarkers, particularly those related to leukocyte composition and sex hormone regulation, could be predicted from MSA data. Despite its reduced CpG coverage, the MSA platform retains sufficient methylation information for biological age and biomarker prediction in the Japanese population.

## List of abbreviations

DNA: Deoxyribonucleic Acid
DNAm: DNA methylation
MSA: Methylation Screening Array
MCV: mean cell volume
RDW-CV: red cell distribution width-coefficient of variation
ALP: alkaline phosphatase
WBC: white blood cell
J-PCPhenoAge: PC-PhenoAge for Japanese Population
ALP/IFCC: ALP based on International Federation of Clinical Chemistry and Laboratory Medicine
ALT: alanine aminotransferase
GPT: glutamic pyruvate transaminase
AST: aspartate aminotransferase
DHEA-S: dehydroepiandrosterone sulfate
FSH: follicle-stimulating hormone
HDL: hematocrit, high-density lipoprotein
Hb: hemoglobin
LD/IFCC: lactate dehydrogenase based on IFCC
LH: luteinizing hormone
MCH: mean corpuscular Hb
MCV: mean corpuscular volume
NT-proBNP: N-terminal pro-brain natriuretic peptide
RBC: red blood cell
WBC: White Blood Cell
γ-GTP: γ-glutamyl transpeptidase
free T3: free triiodothyronine
free T4: free thyroxine
A/G ratio: albumin/globulin ratio
PSA: prostate-specific antigen
MS: mortality score
MAE: mean absolute error
RMSE: root mean squared error
SD: standard deviation

## Declarations

### Ethics approval and consent to participate

This study was conducted in accordance with the Declaration of Helsinki. The study protocol was reviewed and approved by the Asia–Oceania Anti-Aging Promotion Association Ethics Review Committee (approval number: AOAAPA22-001). All participants were informed of the study objectives and procedures, and written informed consent was obtained from all individuals prior to participation.

### Consent for publication

Not applicable.

### Availability of data and materials

The imputed values for all clinical laboratory measurements are provided in Table S2. The original values are available upon reasonable request. Estimated coefficients for each model are provided in Table S3-S6. MAE and RMSE are summarized in Table S7 and S8 respectively. All other data supporting the findings of this study are available from the corresponding author upon reasonable request.

### The Use of Generative AI and AI-assisted Technologies

The authors used ChatGPT to assist with the editing and language during the preparation of this manuscript. The manuscript was edited by the authors using the AI tool. The authors take full responsibility for the content of this manuscript.

## Authors’ contributions

Tatsuma Shoji (TS) performed data analysis, generated all figures and tables, and wrote the manuscript draft. Yui Tomo (YT) provided statistical guidance, contributed to the design of the analytical framework, and reviewed and edited the manuscript draft. Ryo Nakaki (RN) collected the data and led the overall direction and management of the project.

## Acknowledgements

We extend our deepest gratitude to Sawako Hibino, whose efforts in coordinating and collecting blood samples formed the basis of the dataset used in this study. We thank Editage (www.editage.com) for providing English language editing assistance. The contributions of these individuals were indispensable for the successful execution of this study.

## Supplemental Information

Table S1. Proportion of missing values for each of the 59 clinical laboratory measurements.

Table S2. Imputed values for all 59 clinical laboratory measurements across the 214 participants.

Table S3. Estimated coefficients for J-PhenoAge.

Table S4. Estimated coefficients for the MSA-based Phenotypic Age Model.

Table S5. Estimated coefficients for the first-stage model.

Table S6. Estimated coefficients for the second-stage model.

Table S7. MAE between observed and predicted values in Models 1 and 2.

Table S8. RMSE between observed and predicted values in Models 1 and 2.

